# Exosome Trafficking Is a Key Regulator of Adipocyte Thermogenesis

**DOI:** 10.1101/2025.07.20.665768

**Authors:** Devesh Kesharwani, Michele Karolak, Chad Doucette, Rachelle Mendola, Summer Pray, Raghu Bhardwaj, Su Su, Anne Harrington, Victoria DeMambro, Clifford Rosen, Lucy Liaw, Aaron C. Brown

## Abstract

Activation of beige adipocytes enhances energy expenditure and promotes metabolic health, presenting a promising approach for combating obesity and diabetes. As part of this process, thermogenesis, fueled in part by uncoupled mitochondrial respiration, plays a central role in converting calories into thermal energy, thereby preventing their storage as fat. Here, we identify a role for exosome trafficking as an intrinsic regulator of beige adipocyte thermogenesis. Exosomes are small extracellular vesicles that mediate cell-cell and intracellular communication by transporting regulatory cargo, including microRNAs, proteins, and lipids. Using both human cells and mouse models, we show that thermogenic activation of beige adipocytes promotes the rapid release of exosomes enriched in microRNAs known to suppress thermogenic programs. Genetic or pharmacological blockade of exosome secretion attenuates thermogenesis, whereas enhancing exosome release amplifies thermogenic output. Mice deficient in the exosome secretion regulator *Rab27a* exhibit reduced energy expenditure in response to both cold exposure and β3-adrenergic stimulation. These findings establish exosome trafficking as a key contributor to beige adipocyte thermogenic capacity, highlighting an intracellular mechanism that may be leveraged to enhance energy expenditure and treat obesity-related metabolic diseases.

**Significance Statement:** Thermogenic adipocytes, including beige fat cells, help maintain energy balance by converting excess nutrients into heat, thereby reducing fat storage and supporting metabolic function. Although these cells are known to promote energy expenditure, the intracellular processes that enable their full thermogenic response are not well defined. Here, we show that exosome secretion is required for beige adipocytes to reach their full thermogenic potential. Blocking exosome release dampens this response, while boosting it amplifies thermogenic output. These findings point to exosome release as an essential part of thermogenic regulation and a potential target for improving metabolic health.

---

Obesity, defined by excessive lipid accumulation in white adipose tissue, leads to chronic inflammation, insulin resistance, and dysregulated adipokine secretion, which together drive systemic metabolic dysfunction and increase the risk of type 2 diabetes, cardiovascular disease, and certain cancers (1). Lifestyle interventions such as diet and exercise are first-line treatments for obesity, but long-term adherence is often poor and weight regain is common. Although bariatric surgery and GLP-1 receptor agonists are effective treatments for obesity, both are costly, can cause adverse effects, and are frequently associated with weight regain after initial success (2, 3). Given the limitations of current therapies, thermogenic adipocytes, including brown and beige fat cells, have emerged as compelling targets for obesity and diabetes treatment due to their ability to enhance energy expenditure and improve metabolic function through increased lipid and glucose utilization (4–7).

Brown and beige adipocytes play a crucial role in maintaining body temperature and supporting systemic metabolism by burning nutrients to produce heat upon activation (8). Brown adipocytes reside in dedicated depots such as the interscapular, cervical, and spinal regions and are developmentally programmed for stable thermogenic activity. In contrast, beige adipocytes are inducible thermogenic cells within subcutaneous white fat that acquire reversible thermogenic activity in response to changes in temperature (8, 9). Upon cold exposure, norepinephrine released from sympathetic nerves activates brown and beige adipocytes through *α*1- and *β*3-adrenergic signaling (10, 11). Both brown and beige adipocytes contain multilocular lipid droplets, are rich in mitochondria, and express uncoupling protein-1 (UCP1), which drives heat generation by uncoupling mitochondrial respiration (4). In addition to UCP1-dependent thermogenesis, several UCP1-independent energy-dissipating mechanisms, including calcium flux, creatine turnover, and fatty acid cycling, consume ATP without driving cellular activity and release that energy as heat (9). Recent studies indicate that beige adipocytes can be subdivided into distinct populations based on their reliance on these various pathways (12, 13).

Upon cold exposure or *β*-adrenergic activation, brown and beige adipocytes enhance glucose uptake and fatty acid oxidation to fuel thermogenesis, a process that supports thermal homeostasis and also improves insulin sensitivity, facilitates triglyceride clearance, and contributes to systemic glucose regulation (4, 6). The discovery of beige fat and its inducible nature has sparked interest in its therapeutic potential, with efforts underway to pharmacologically recruit subcutaneous beige fat to combat obesity and metabolic disease. However, current approaches are limited: *β*-adrenergic receptor agonists can carry cardiovascular risks, while prolonged cold exposure is often impractical and may pose safety concerns for individuals with elevated cardiovascular risk (14). These challenges underscore the need for alternative strategies to safely and effectively harness thermogenic adipocytes for metabolic benefit.

Thermogenic adipocytes also act as endocrine cells that influence distant organs and systemic metabolism by secreting adipokines and other bioactive factors, including lipids and peptides, which help regulate glucose homeostasis, lipid metabolism, and insulin sensitivity (15, 16). Among these secreted factors, exosomes, small extracellular vesicles that carry microRNAs, proteins, lipids, and RNAs, have emerged as important mediators of intercellular communication with systemic metabolic effects (17).

Adipose-derived exosomes can exert both beneficial and detrimental effects depending on physiological state and tissue origin. In obesity, white adipose tissue-derived exosomes are often enriched with proinflammatory cargo that promotes insulin resistance and metabolic dysfunction (17). However, they may also serve adaptive functions. For example, exosomes from adipose tissue of obese mice deliver proteins to pancreatic *β*-cells, enhancing insulin secretion and glucose control, suggesting they signal insulin resistance in adipose tissue to boost insulin production when needed (18). In contrast, exosomes from thermogenic depots support metabolic homeostasis. Brown adipose tissue-derived exosomal miR-132-3p, regulated by norepinephrine during cold exposure, inhibits hepatic lipogenesis by targeting *Srebf1*, thereby facilitating the body’s adaptation to cold stress (19). Similarly, cold exposure induces secretion of miR-99b-5p-enriched exosomes from brown fat that target the liver to regulate expression of fibroblast growth factor 21 (FGF21), a key metabolic hormone involved in energy expenditure and glucose homeostasis (20). Circulating levels of exosomal miR-92a inversely correlate with brown fat activity in humans, suggesting its potential use as a non-invasive biomarker of thermogenic capacity (21). Exosomes from adipose tissue macrophages of lean mice also improve glucose tolerance and insulin sensitivity when administered to obese recipients (21). While these findings highlight the complex endocrine roles of adipose-derived exosomes, it remains unclear whether exosome secretion also plays a direct, cell-autonomous role in thermogenic adipocyte activation.

Here, we show that beige adipocyte activation is associated with increased exosome release containing suppressive cargo, leading us to hypothesize that exosome secretion is linked to thermogenesis. Supporting this, inhibition of exosome secretion reduced UCP1 expression and thermogenic output, whereas enhanced secretion boosted thermogenesis. Loss of the exosome trafficking protein RAB27A diminished exosome release and impaired thermogenic capacity, resulting in reduced energy expenditure in response to cold or *β*_3_-adrenergic stimulation. These findings identify exosome secretion as a cell-intrinsic mechanism that facilitates beige adipocyte thermogenesis and supports systemic energy balance.

## Results

### Exosome release during beige adipocyte activation is associated with thermogenic potential

In adult mammals, the generation of beige adipocytes from adipose-resident precursor cells of stromal, perivascular, and smooth muscle origin contributes modestly to overall beige adipocyte mass and occurs primarily after prolonged cold exposure (22). In contrast, most beige adipocytes arise in response to acute cold exposure via rapid reprogramming of “whitened” beige adipocytes into a brown-like state (browning), which is defined by acquisition of markers of thermogenic adipocytes, including UCP1 (23). To examine this adaptive process in greater detail, we employed a human iPSC-derived beige adipocyte model to investigate exosome release during browning of mature adipocytes and to characterize their microRNA cargo (24). iPSC-derived beige adipocyte precursors were first differentiated into mature beige adipocytes over 12 days using a defined adipogenic cocktail (24). These mature beige adipocytes were then cultured in non-adipogenic basal medium for 4 days to induce a “whitening” process, which resulted in a marked reduction in *UCP1* expression (24). To mimic cold-induced browning, whitened cells were treated for 6 hours with forskolin, a *β*-adrenergic analog that activates adenylyl cyclase to elevate intracellular cAMP. This short-term stimulation increased the expression of *UCP1* and *PGC1α*, markers of mitochondrial uncoupling and biogenesis, respectively (Fig. 1A). Nanoparticle tracking analysis revealed that forskolin activation triggered an *>*8-fold increase in exosome release into the culture medium (Fig. 1B–C). Based on published literature, we compiled a panel of 98 microRNAs previously linked to adipogenesis, metabolism, and obesity (Fig. S1A) (25–29). We then developed targeted qPCR assays to quantify these microRNAs in exosomes secreted by beige adipocytes under both whitened and forskolin-stimulated conditions (Fig. 1D, *P <* 0.02; see Fig. S1A for the full data set).

**Fig. 1.**
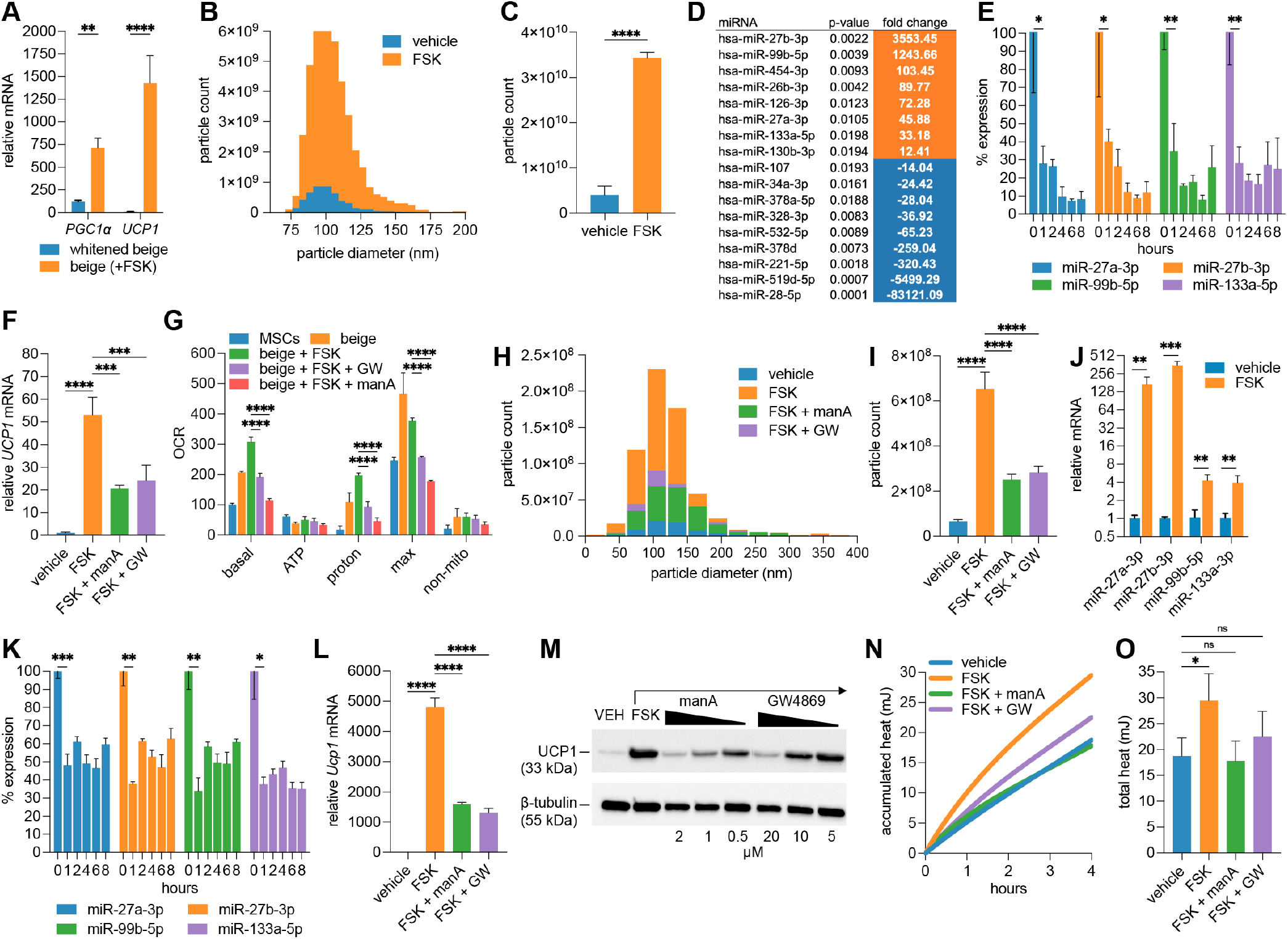
Exosome secretion correlates with thermogenic activation in beige adipocytes and is conserved across human and mouse models. (A) RT-qPCR analysis of *PGC1α* and *UCP1* transcript levels in human iPSC-derived beige adipocytes following 6-hour forskolin treatment (10 *µ*M, *n* = 3). (B, C) Nanoparticle tracking analysis (NTA) of exosomes from beige adipocytes ± forskolin treatment for 6 hours, showing (B) size distribution and (C) *>*8-fold increase in concentration (*n* = 3). (D) Differentially expressed microRNAs in exosomes from beige adipocytes with or without forskolin treatment (*n* = 3), ranked by fold change. Orange indicates increased expression with forskolin, blue indicates decreased expression. (E) Time-course qPCR analysis of intracellular miR-27a/b-3p, miR-99b-5p, and miR-133a-5p in beige adipocytes following forskolin treatment, showing a marked reduction by 1 h (*n* = 3 per time point). Time zero normalized to 100%. (F) *UCP1* expression in forskolin-treated beige adipocytes is reduced by co-treatment with exosome biogenesis and release inhibitors manumycin A (ManA) or GW4869 (GW) (*n* = 3). (G) Oxygen consumption rate (OCR) analysis via Seahorse XF analyzer showing reduced proton leak-linked respiration in forskolin-stimulated beige adipocytes ± exosome inhibitors (*n* = 3). (H, I) NTA of exosomes from mouse beige adipocytes ± forskolin and inhibitors: (H) size distribution by bin; (I) total particle concentration (*n* = 5). (J, K) qPCR showing forskolin-induced enrichment of miR-27a/b, miR-99b-5p, and miR-133a-5p in mouse beige-derived exosomes after 4-hour treatment (J) and their corresponding intracellular depletion (K) (*n* = 3). (L) *Ucp1* mRNA expression in forskolin-treated mouse beige adipocytes ± exosome inhibitors after 6-hour treatment (*n* = 3). (M) Western blot showing UCP1 protein reduction in a dose-dependent manner with exosome inhibitor co-treatment (*n* = 3 pooled replicates with *β*-tubulin as loading control). (N, O) Microcalorimetry of cultured beige adipocytes showing (N) average heat accumulation over time and (O) total heat output (quantified from N, with error bars) in the presence or absence of forskolin and exosome inhibitors (*n* = 3). Refer to Methods for statistical tests and significance thresholds.

Several microRNAs were significantly up- or downregulated in exosomes following beige adipocyte browning, suggesting that thermogenic activation alters the selective packaging and secretion of microRNAs. Among the most enriched was miR-99b-5p (*>*1200-fold), consistent with prior reports showing its secretion from brown adipose tissue (BAT) following cold exposure and its role in targeting *Fgf21* in the liver (Fig. 1D) (30). FGF21 is an adipokine secreted by both beige and brown adipocytes and plays a key role in promoting browning and enhancing thermogenesis in white fat (31). The most highly enriched microRNA in exosomes from beige adipocytes undergoing browning was miR-27b-3p (∼3500-fold). Its paralog miR-27a-3p, which shares an identical seed sequence, was also significantly enriched (∼46-fold), consistent with coordinated regulation within the miR-27 family. Previous studies have shown that cellular levels of miR-27a and miR-27b are downregulated during beige and brown adipocyte differentiation *in vitro*, as well as in subcutaneous and brown adipose depots following acute cold exposure (32). The miR-27 family targets and suppresses translation of multiple transcription factors critical for UCP1-dependent thermogenesis, and *in vivo* knockdown of miR-27b enhances browning and improves metabolic outcomes in obese mice (32, 33). Loss of miR-27, therefore, is predicted to be required for the thermogenic adipocyte phenotype. Other microRNAs enriched in exosomes from forskolin-activated beige adipocytes included known inhibitors of adipogenesis (miR-130b-3p, miR-454-3p), thermogenesis (miR-133a-5p), and insulin signaling (miR-126-3p) (34–37). Cellular levels of thermogenesis-associated microRNAs (miR-99b-5p, miR-27a/b-3p, and miR-133a-5p) were found to decrease rapidly (within 1 hour) following forskolin stimulation, consistent with their selective export during browning (Fig. 1E). To test whether this exosome secretion impacts thermogenesis, beige adipocytes were stimulated with forskolin in the presence of the exosome secretion inhibitors manumycin A or GW4869 (38, 39). Both inhibitors partially blocked forskolin-induced *Ucp1* expression (Fig. 1F). Furthermore, oxygen consumption assays revealed that proton leak-linked respiration was suppressed in the presence of manumycin A or GW4869, indicating reduced mitochondrial uncoupling and thermogenic activity (Fig. 1G). Together, these findings are consistent with a role for exosome secretion in beige adipocyte thermogenesis, potentially by promoting the export of anti-thermogenic microRNAs or other exosomal cargo that may influence thermogenic regulation.

### Exosome-associated thermogenesis is conserved between humans and mice

Similar to human cells, whitened beige adipocytes generated from the stromal vascular fraction of mouse inguinal adipose tissue exhibited a substantial increase in exosome secretion following 6 hours of forskolin treatment, which was reversed by co-treatment with the exosome secretion inhibitors manumycin A and GW4869 (Fig. 1H-I). qPCR analysis of exosomal RNA revealed enrichment of miR-27a/b, miR-99b-5p, and miR-133a-5p in exosomes from forskolin-activated beige adipocytes (Fig. 1J), accompanied by a corresponding decrease in their intracellular levels (Fig. 1K). Inhibition of exosome secretion with manumycin A and GW4869 partially attenuated the forskolin-induced upregulation of *Ucp1* transcript levels during beige adipocyte browning (Fig. 1L) and suppressed UCP1 protein levels in a dose-dependent manner (Fig. 1M). Comparable results were observed when beige adipocytes were activated with cAMP instead of forskolin, indicating that the effects were not due to off-target actions of this plant-derived adenylyl cyclase activator (Fig. S1B). Furthermore, exosome secretion inhibition significantly reduced heat production during beige adipocyte browning, as measured by high-resolution microcalorimetry (Fig. 1N-O). These results indicate that the role of exosome secretion in beige adipocyte thermogenesis is conserved across species and that mouse models can provide valuable insight into exosome-regulated browning processes relevant to human physiology.

### Genetic enhancement of exosome secretion increases beige adipocyte thermogenesis

To determine whether increasing exosome secretion enhances beige adipocyte thermogenesis, we generated transgenic doxycycline-inducible mice expressing an exosome secretion booster cassette (Tg-ExoBooster; see schematic in Fig. 2A). This cassette includes the genes *STEAP3, SDC4*, and *NadB*, which promote exosome biogenesis and multivesicular body formation, thereby enhancing exosome release (40). Following 72 hours of doxycycline treatment in both wild-type and Tg-ExoBooster inguinal SVF-derived beige adipocytes, we confirmed transgene expression by RT-PCR (Fig. 2B) and observed a ∼ 13-fold increase in exosome secretion in the transgenic group (Fig. 2C–D). This increase was partially reversed by co-treatment with exosome secretion inhibitors manumycin A and GW4869, suggesting pathway overlap (Fig. 2D). Beige adipocytes expressing the ExoBooster cassette exhibited elevated *Ucp1* transcript levels and increased UCP1 protein following forskolin stimulation compared to wild-type controls, at 6 (RNA) and 24 (protein) hours following treatment, indicating a hypersensitized thermogenic response (Fig. 2E-F). Furthermore, Tg-ExoBooster-positive beige adipocytes maintained higher *Ucp1* expression after whitening in non-adipogenic basal medium compared to wild-type adipocytes (Fig. S2A). These cells also produced significantly more heat than wild-type beige adipocytes, as measured by microcalorimetry (Fig. 2G-H). To test whether this enhanced thermogenic phenotype was retained *in vivo*, Tg-ExoBooster mice were housed at thermoneutrality (29C) for 7 days to promote whitening of adipose tissue and stabilize basal energy expenditure, treated with doxycycline from days 2–7, and then subjected to a 3-day cold challenge. Inguinal adipose tissue isolated from Tg-ExoBooster mice exhibited elevated *Ucp1* expression (Fig. 2I) and generated more heat *ex vivo* than tissue from wild-type mice (Fig. 2J-K), as determined by microcalorimetry. In contrast, no significant changes in *Ucp1* expression or heat production were observed in brown adipose tissue under the same temperature conditions (Fig. S2B–D). Collectively, these data demonstrate that genetic enhancement of exosome secretion can promote beige adipocyte thermogenesis both *in vitro* and *in vivo*.

**Fig. 2.**
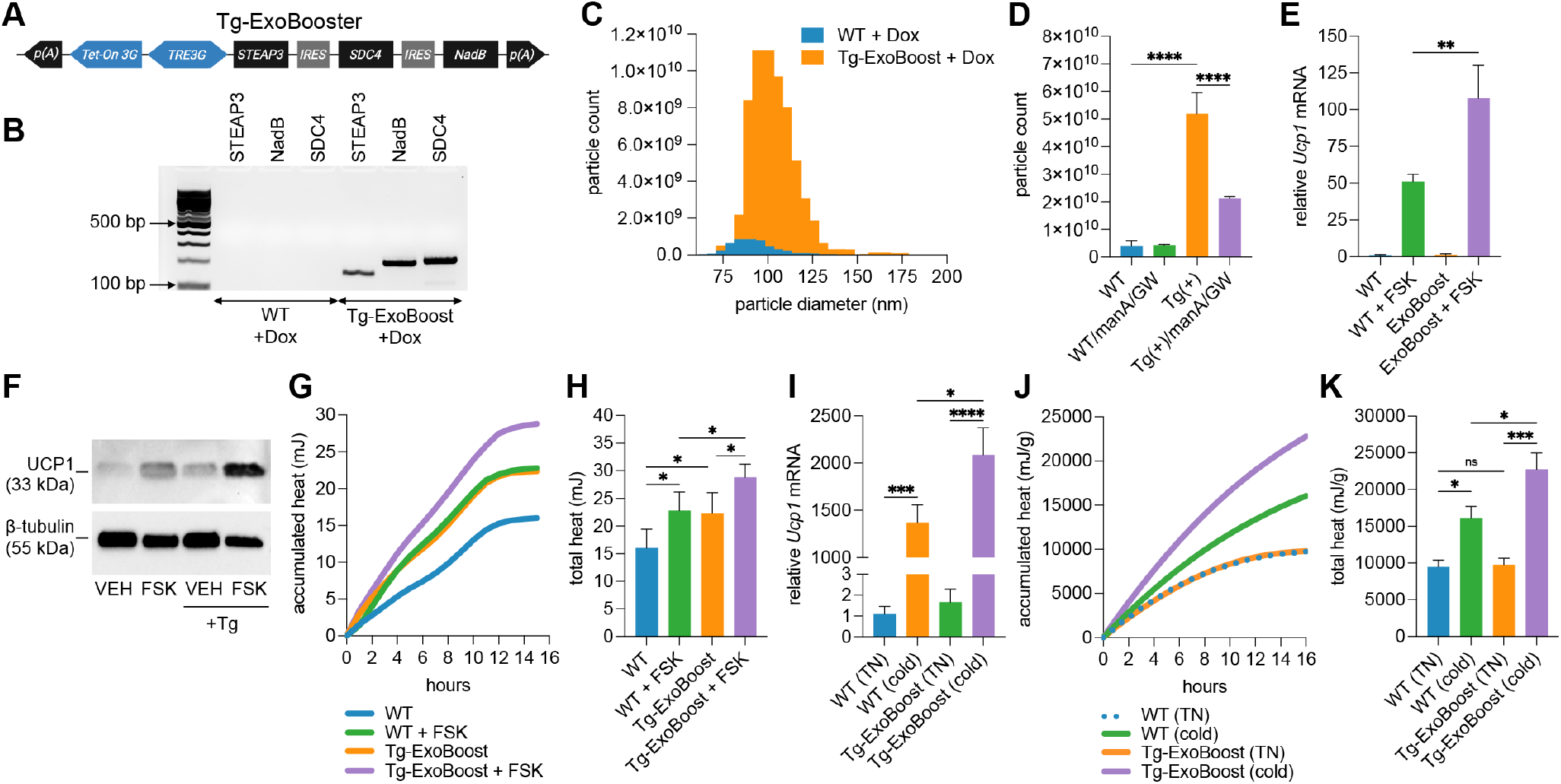
Genetically boosted exosome secretion enhances thermogenesis in beige adipocytes. (A) Schematic of the Tg-ExoBooster construct. The bidirectional TRE3G promoter drives upstream expression of Tet-On 3G and downstream expression of a tricistronic STEAP3–IRES–SDC4–IRES–NadB cassette after addition of doxycycline. p(A), polyadenylation signal; IRES, internal ribosome entry site. (B) RT-PCR validation of ExoBooster transgene expression in cultured beige adipocytes after 72 hours doxycycline treatment (*n* = 3). (C) Nanoparticle tracking analysis (NTA) of exosome size distribution from Tg-ExoBooster versus wild-type (WT) beige adipocytes treated with doxycycline. NTA analysis of total exosome particle counts from Tg-ExoBooster versus WT beige adipocytes treated with doxycycline and with or without inhibitor co-treatment (*n* = 3). qPCR analysis of *Ucp1* expression in Tg-ExoBooster and WT beige adipocytes after 6-hour forskolin stimulation (*n* = 3). (F) Western blot analysis of UCP1 protein levels from pooled replicates (*n* = 3). (G, H) Microcalorimetry traces showing (G) average heat accumulation over time and (H) total heat output (quantified from G, with error bars) in forskolin-treated Tg-ExoBooster and wild-type beige adipocytes following doxycycline treatment (*n* = 5). (I) RT-qPCR analysis of *Ucp1* expression in inguinal adipose tissue from wild-type and Tg-ExoBooster mice after 3 days doxycycline feed and cold challenge (*n* = 5). (J, K) *Ex vivo* microcalorimetry traces (J) and quantified heat output (K) from inguinal fat pads post-doxycycline and cold challenge (*n* = 5). Heat data in J–K normalized to tissue weight in grams (g).

### RAB27A deficiency impairs exosome secretion and ther-mogenesis in beige adipocytes

RAB27A is critical for the fusion of multivesicular bodies with the plasma membrane and the subsequent release of exosomes (41). To determine whether impaired exosome secretion compromises thermogenic function, we used a global *Rab27a* null mouse model (42). Beige adipocytes were differentiated from the stromal vascular fraction of inguinal adipose tissue and then exposed to either forskolin or vehicle to assess their thermogenic capacity. Exosomes were isolated from the conditioned media and analyzed for particle number. Both *Rab27a* heterozygous and null beige adipocytes secreted significantly fewer exosomes than wild-type cells under both vehicle- and forskolin-stimulated conditions (Fig. 3A-B), confirming a substantial impairment in exosome release. A loss of thermogenic function accompanied this defect. *In vitro, Rab27a* heterozygous and null beige adipocytes showed markedly reduced forskolin-induced *Ucp1* mRNA expression and UCP1 protein levels compared to wild-type controls (Fig. 3C–D), along with significantly diminished forskolin-stimulated heat production (Fig. 3E–F). To assess whether these deficits extended *in vivo*, wild-type and *Rab27a* null mice were housed at thermoneutrality (29C) for 7 days to normalize the whitening of adipose tissue, followed by a 7-day challenge of cold exposure. Inguinal adipose tissue from cold-exposed wild-type mice exhibited robust induction of *Ucp1* transcription and UCP1 protein expression, whereas *Rab27a* null mice showed reduced responses, with *Ucp1* mRNA and protein levels decreased by approximately 60% and 40%, respectively, compared to wild-type controls (Fig. 3G–H; Fig. S3). Microcalorimetry revealed inguinal fat pads from cold-treated wild-type mice generated significantly more heat than those from *Rab27a* null mice (Fig. 3I–J). These experiments support the conclusion that RAB27A-dependent exosome trafficking is a critical component of the thermogenic signaling cascade and that disruption of this pathway compromises the ability of adipose tissue to properly respond to thermogenic cues. The observation that *Rab27a* heterozygous mice consistently phenocopied nulls across all assays suggests that beige adipocyte thermogenesis may require a threshold level of RAB27A function, raising the possibility of haploinsufficiency or other dosage-sensitive mechanisms related to exosome secretion.

**Fig. 3.**
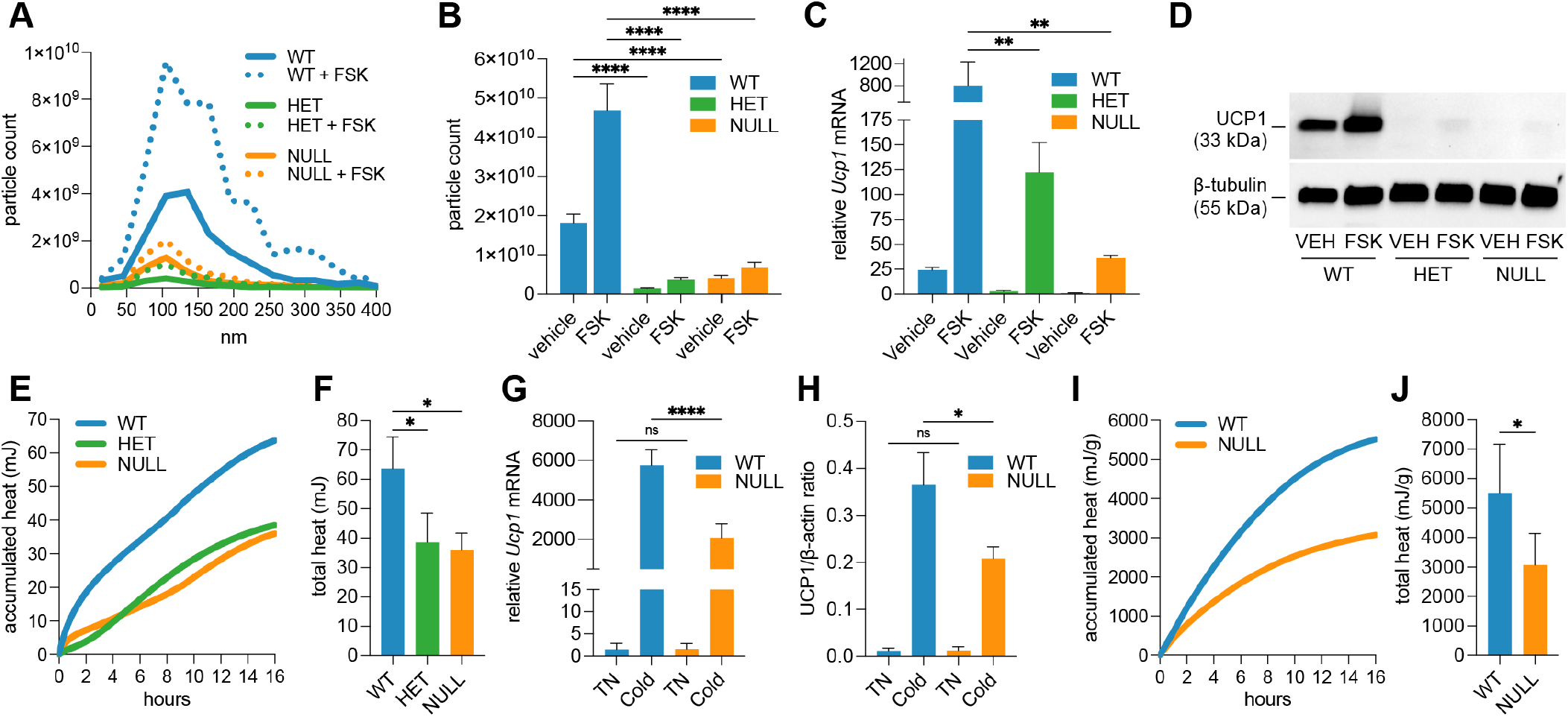
*Rab27a* deficiency impairs exosome secretion and thermogenesis in beige adipocytes. (A, B) Nanoparticle tracking analysis showing (A) average exosome count and size distribution and (B) total exosome particle counts (quantified from A, with error bars) from wild-type, *Rab27a* heterozygous, and *Rab27a* null beige adipocytes ± forskolin (*n* = 4). (C) qPCR analysis of *Ucp1* mRNA expression in cultured beige adipocytes ± forskolin (*n* = 3). (D) Western blot of UCP1 protein in cultured beige adipocytes (*n* = 3 replicates pooled per treatment). *β*-tubulin serves as loading control. (E, F) Microcalorimetry traces showing (E) average heat accumulation over time and (F) total heat output (quantified from E, with error bars) from beige adipocytes (*n* = 3). (G) *Ucp1* expression in inguinal fat following cold exposure in wild-type, *Rab27a* heterozygous, and null mice (*n* = 5 mice per group). (H) Immunoblot quantitation of UCP1 protein from inguinal adipose tissue (*n* = 6 mice per group). Data are normalized to *β*-tubulin (see Fig. S3A for blot). (I, J) Microcalorimetry traces showing (I) average heat accumulation over time and (J) total heat output (quantified from I, with error bars) from inguinal adipose tissue of RT and cold-exposed mice (*n* = 4 per group). Heat data in I–J normalized to tissue weight (g).

### RAB27A deficiency reduces energy expenditure in response to cold or *β*_3_-adrenergic stimulation

To determine whether *Rab27a* deficiency affects whole-body metabolism, wild-type and *Rab27a* heterozygous mice were housed at room temperature (∼ 22^*°*^C) or exposed to cold (6^*°*^C) for 7 days, then transferred to metabolic cages at room temperature. After a 24-hour acclimation period, energy expenditure, respiratory exchange ratio, physical activity, food intake, and water consumption were measured over the next 48 hours (see Supplementary Table 1 for detailed measurements and corresponding p-values). Body weight was similar between wild-type and *Rab27a* heterozygous mice under both room temperature (RT) and cold conditions; however, heterozygous mice displayed significantly higher fat mass at RT (*p* = 0.0005), as shown in Fig. 4A. This excess fat was lost during cold exposure, resulting in comparable fat mass across genotypes by day 7. Indirect calorimetry revealed no difference in total energy expenditure between genotypes at RT over the 48-hour recording period (Fig. 4B). However, after 7 days of cold exposure, *Rab27a* heterozygous mice exhibited reduced energy expenditure compared to wild-type controls (Fig. 4C). Quantification of total 24-hour energy expenditure showed a significant reduction in heterozygous mice under cold conditions (*p* = 0.02; Fig. 4D), with the most pronounced difference observed during the daytime phase (*p* = 0.0037; Fig. 4E). A similar trend was observed during the nighttime phase, although the difference did not reach statistical significance (*p* = 0.0993; Fig. 4F). Further analysis of resting energy expenditure (REE) revealed significantly lower values in heterozygous mice during both daytime (*p* = 0.0362; Fig. 4G) and nighttime (*p* = 0.0149; Fig. 4H) cold exposure periods. The combined 24-hour REE was also significantly reduced in *Rab27a* heterozygous mice compared to wild-type (*p* = 0.0114; Fig. 4I), indicating a deficit in basal thermogenic output. Ambulatory activity, assessed by total distance walked in the cage, was significantly reduced in *Rab27a* heterozygous mice under RT conditions (*p* = 0.0462; Fig. 4J), though this difference did not persist under cold exposure. No significant differences were observed in respiratory exchange ratio (RER), food or water intake, or sleep duration between genotypes under either temperature condition (see Supplementary Table 1). In combination, these data indicate that *Rab27a* heterozygosity impairs cold-induced increases in resting and total energy expenditure, independent of behavioral or nutritional factors, supporting a role for *Rab27a* in the regulation of basal thermogenesis.

**Fig. 4.**
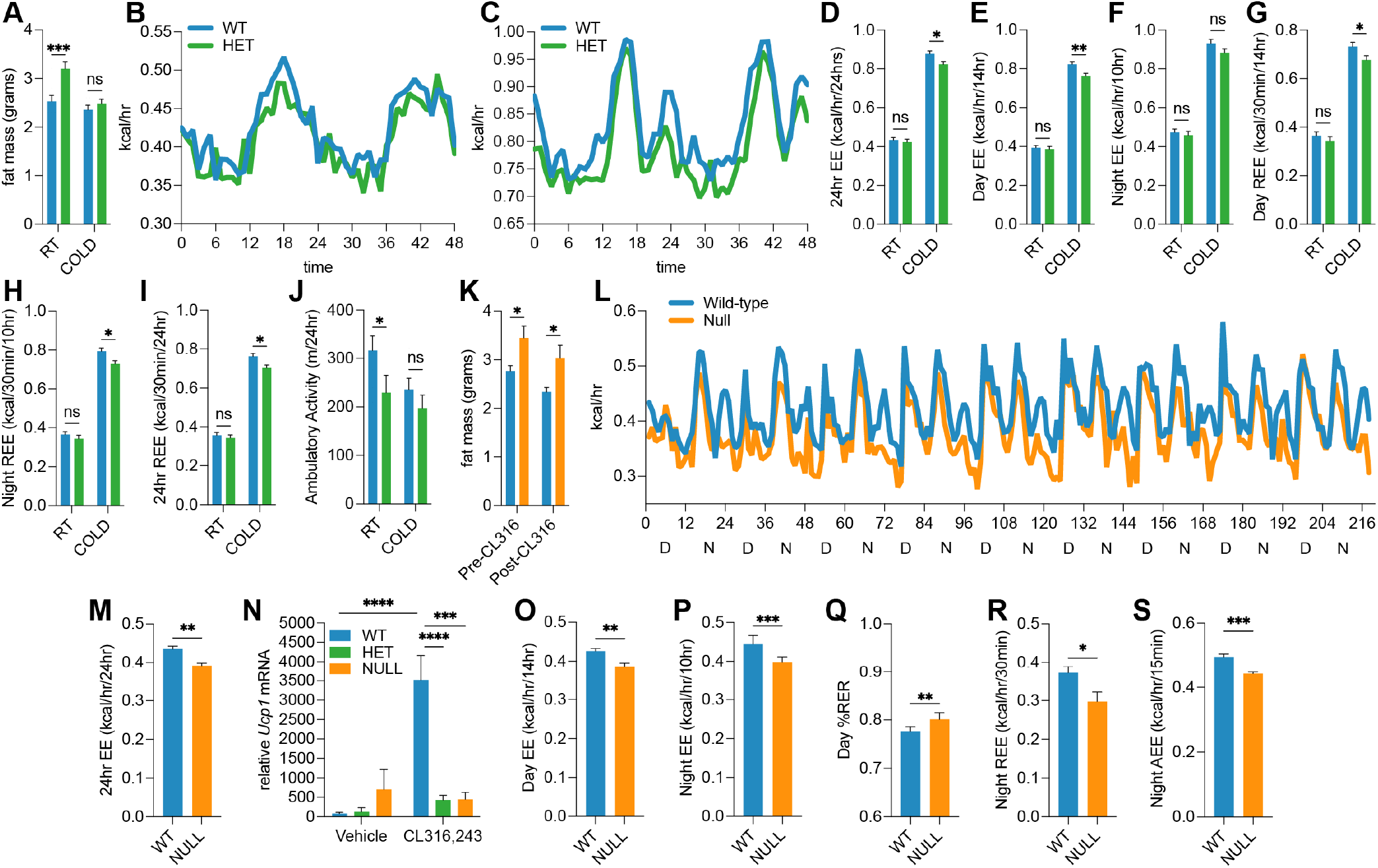
*Rab27a* deficiency reduces energy expenditure in response to cold and *β*_3_ -adrenergic stimulation. (A) Fat mass of wild-type and *Rab27a* heterozygous female mice housed at room temperature (RT, ∼22^*°*^C) or cold exposed (6^*°*^C) for 7 days (*n* = 6–8 mice per group for A–J). (B, C) Metabolic cage analysis of energy expenditure over time at (B) room temperature and (C) following cold exposure. (D–F) 24-hour, daytime, and nighttime energy expenditure after cold exposure. (G–I) Daytime, nighttime, and combined 24-hour resting energy expenditure (REE). (J) Ambulatory activity at RT and cold. (K) Fat mass before and after CL316,243 treatment in wild-type and *Rab27a* null mice (*n* = 6 mice per group for L–S). (L) Average total energy expenditure during 9-day CL316,243 treatment. (M) Average 24-hour energy expenditure over 9 days. (N) RT-qPCR analysis of *Ucp1* mRNA in inguinal adipose tissue *±* CL316,243 treatment (day 9). (O, P) Average daytime and nighttime energy expenditure, (Q) daytime RER, and (R) active energy expenditure (AEE) over 9 days. Refer to Methods for statistical tests and significance thresholds.

To assess the role of *Rab27a* in *β*_3_-adrenergic-driven thermogenesis, wild-type and *Rab27a* null mice were treated with CL316,243 (1 mg/kg) daily for 9 days during metabolic cage analysis (see Supplementary Table 2 for complete data). Untreated controls were excluded, as wild-type baselines were established in the cold exposure study, and *Rab27a* heterozygous mice displayed similar phenotypes to nulls (Fig. 3A–F). As with the cold study, at baseline, *Rab27a* null mice had significantly greater total fat mass (in grams) than wild-type controls (*p* = 0.0270; Fig. 4K; *n* = 8 mice per group). Although both groups lost similar proportions of fat mass (12–15%) after treatment, wild-type mice were ∼4% heavier by the end of the study (*p* = 0.0323; Supplementary Table 2). This increased body weight in wild-type mice appears to result from a significant increase in lean mass (+6%, *p* = 0.0087), which the null mice failed to achieve (*−* 0.1%; *p* = 0.0051 compared to WT); a known outcome of CL316,243-induced anabolic remodeling of skeletal muscle in rodents (43, 44). Total energy expenditure over the 9-day period was reduced in null mice (Fig. 4L), and average 24-hour energy expenditure across this period was significantly lower in nulls (*p* = 0.0020; Fig. 4M). Statistical comparisons for individual day and nighttime intervals are shown in Fig. S3B. Similar to the effects observed under cold exposure (Fig. 3G), both *Rab27a* heterozygous and null mice exhibited significantly lower *Ucp1* transcript levels following CL316,243 treatment compared to wild-type controls (Fig. 4N), consistent with a blunted thermogenic response. Both average daytime (*p* = 0.0062) and nighttime (*p* = 0.0014) energy expenditure were reduced in *Rab27a* null mice (Fig. 4O and 4P). Respiratory exchange ratio (RER) was significantly elevated during the daytime in *Rab27a* null mice (*p* = 0.0056; Fig. 4Q), suggesting a shift toward carbohydrate metabolism and reduced fatty acid oxidation, possibly due to lower UCP1 expression in the nulls (Fig. 3G and H). Resting energy expenditure (REE) was also broadly impaired. Both nighttime REE (*p* = 0.0326; Fig. 4R) and 24-hour REE (*p* = 0.0012, Supplementary Table 2) were significantly reduced in null mice, indicating a persistent basal thermogenic defect. Nighttime active energy expenditure (AEE) was also lower in *Rab27a* null mice (*p* = 0.0003; Fig. 4S), compounding the reduction in total energy output. No significant differences were observed in food intake, water consumption, ambulatory activity (beam breaks, walking distance), or sleep duration (Supplementary Table 2), excluding behavioral confounders.

Taken as a whole, these experiments reveal that RAB27A is required for the full thermogenic and anabolic response to both cold and *β*_3_-adrenergic stimulation. In its absence, mice exhibit reduced energy expenditure, impaired lean mass gain, and altered substrate utilization, highlighting the critical role of RAB27A and exosome secretion in adaptive thermogenesis.

## Discussion

This study uncovers a previously unappreciated role for exosome secretion in supporting the thermogenic capacity of beige adipocytes. We show that activation of beige adipocytes is associated with increased exosome release and that these exosomes are enriched in microRNAs typically linked to the suppression of thermogenic programming. Impairing exosome secretion, either through *Rab27a* deficiency or pharmacologic inhibition with the small molecules manumycin A and GW4869, leads to a blunted thermogenic response. Mice lacking *Rab27a* exhibit significantly reduced energy expenditure in response to cold exposure and *β*_3_-adrenergic stimulation, underscoring the functional importance of this pathway *in vivo*. Conversely, genetic enhancement of exosome secretion via a multi-gene booster cassette (ExoBooster) increases UCP1 expression and heat output from beige adipocytes. In sum, these results establish that exosome secretion is not simply a byproduct of adipocyte browning but rather a necessary component of the thermogenic process. Our findings align with and extend the established concept that exosomes are critical mediators of intercellular communication, particularly in metabolic regulation (17). We propose a model in which beige adipocytes actively secrete exosomes enriched in inhibitory microRNAs as a potential self-regulatory mechanism to sustain thermogenic activation. Regulation of these intracellular microRNAs may be necessary to de-repress thermogenic transcription factors and enable full activation of mitochondrial pathways. This model is supported by the observation that inhibition of exosome secretion blunts forskolin-induced UCP1 expression and thermogenesis, and that *Rab27a*-deficient adipocytes exhibit impaired UCP1 induction, reduced heat production, and altered substrate utilization. Furthermore, the elevation in RER in *Rab27a*-null mice during *β*_3_-adrenergic stimulation (Fig. 4Q) suggests a shift toward carbohydrate metabolism and impaired fatty acid oxidation, possibly due to reduced UCP1 expression (Fig. 3).

While our study primarily focuses on beige adipocytes, exosome-mediated regulation may also occur in classical brown fat and other adipose depots, including visceral fat. Additional studies are needed to determine whether this pathway functions similarly across these distinct adipose tissues. However, exosome secretion may play a more prominent role in beige adipocytes, which must transition from a relatively quiescent or “whitened” state back to a thermogenically active phenotype. This process likely requires more extensive cellular remodeling and may depend on additional regulatory layers, including mechanisms for clearing inhibitory factors during browning. Moreover, although our findings center on UCP1-dependent thermogenesis, it remains possible that exosome secretion also influences UCP1-independent pathways, such as futile cycling of calcium, creatine, or lipids (9). Given the emerging heterogeneity among beige adipocytes, it is also plausible that only specific subsets rely on exosome secretion for full thermogenic activation, an area warranting further investigation.

Importantly, our findings have clinical relevance for obesity and type 2 diabetes. Beige and brown adipocytes are well-established targets for increasing energy expenditure and improving glucose metabolism (4, 7). Strategies to enhance their activity have traditionally relied on cold exposure or *β*_3_-adrenergic agonists, both of which have translational limitations (14). Our work identifies exosome secretion as a novel and targetable mechanism to augment thermogenesis in beige adipocytes, potentially offering a new approach for increasing energy expenditure in obese individuals. Furthermore, because *Rab27a*-deficient mice fail to increase lean mass in response to CL316,243, this work raises the possibility that exosome pathways also intersect with anabolic remodeling and muscle–adipose crosstalk (43, 44).

There are, however, several limitations to consider. First, while our data support a functional role for exosome secretion in promoting thermogenesis, it remains unclear whether this effect is mediated by specific exosomal cargo or by broader aspects of the secretion process itself. Our candidate screen identified several microRNAs (miR-27a/b, miR-133a, miR-99b) that are typically linked to repression of thermogenic gene programs (30, 32, 34), but further studies are needed to determine whether their export plays an active role in adipocyte browning or simply correlates with it. Second, the global *Rab27a* knockout model has known effects beyond adipose tissue (42), which may confound some systemic phenotypes. Future work using adipocyte-specific *Rab27a* deletion models would help isolate the cell-autonomous effects. Lastly, while we demonstrate effects in mouse and iPSC-derived human models (24), further work is required to confirm the relevance of this pathway in human obesity.

In summary, this study reveals a novel regulatory axis linking exosome secretion to beige adipocyte thermogenesis. By demonstrating that both enhancement and disruption of exosome pathways directly impact energy expenditure, this work provides new insight into adipocyte biology and identifies exosome secretion as a potential target for antiobesity therapies.

## ACKNOWLEDGMENTS

This work was supported by NIH COBRE award P20GM121301 (A. Brown, L. Liaw, and C.J. Rosen) and NIDDK award 1R01DK124261 (A. Brown). The project utilized the services of Core Facilities funded by NIH award U54GM115516 (C.J. Rosen, PI). We kindly thank Drs. Daniel Bojar and Martin Fussenegger for providing the exosome booster plasmids.

## Supporting Information Appendix (SI)

The Supporting Information (SI Appendix) includes additional figures and tables that complement the data presented in the main text.

## Materials and Methods

### Statistical Analysis

Statistical Analysis. Unless otherwise stated, all data are presented as mean ± standard deviation (SD) for *in vitro* experiments and mean ± standard error of the mean (SEM) for *in vivo* studies. The number of biological replicates (n) is indicated in the figure legends. Comparisons between two groups were analyzed using unpaired two-tailed Student’s *t*-tests. For multiple group comparisons, one-way or two-way analysis of variance (ANOVA) was performed as appropriate, followed by Tukey’s multiple comparisons test. MicroRNA profiling statistics (Fig. 1D) were performed using Global Pattern Recognition software (45). A *p*-value *<* 0.05 was considered statistically significant. Significance levels are indicated as follows: *P <* 0.05 (*), *P <* 0.01 (**), *P <* 0.001 (***), *P <* 0.0001 (****).

### Differentiation and Activation of Human and Murine Beige Adipocytes

Human iPSC-derived beige adipocytes were differentiated for 12 days using an adipogenic induction cocktail in EGM2 medium (Lonza), as previously described (24). To induce whitening, mature beige adipocytes were cultured for 4 days in non-adipogenic basal medium consisting of high-glucose DMEM (ATCC, 30-2002) supplemented with 10% exosome-depleted fetal bovine serum (System Biosciences, EXO-FBS-250A-1). Where indicated, cells were treated with 2 *µ*M manumycin A (Sigma, M6418) or 10 *µ*M GW4869 (Sigma, D1692) for 2 hours at the whitening phase to inhibit exosome secretion. Murine beige adipocytes were derived from the stromal vascular fraction (SVF) of inguinal white adipose tissue harvested from female 8-week-old C57BL/6J mice. Tissue was finely minced with a sterile razor blade and digested in isolation buffer containing 0.1% collagenase P (Roche) at 37^*°*^C for 45– 60 minutes on a nutator, with brief vortexing every 10 minutes. The digest was filtered through a 70 *µ*m cell strainer and centrifuged at 300 × *g* for 5 minutes. The SVF pellet was washed with autoMACS running buffer (Miltenyi), centrifuged again, and resuspended in primary culture medium (high-glucose DMEM + 20% FBS + 1% penicillin–streptomycin). Passage 2–3 cells were differentiated for 6 days using the same human adipogenic induction cocktail in DMEM + 10% FBS (ATCC, 30-2020), followed by 6 days of whitening in DMEM + 10% exosome-depleted FBS. Thermogenic activation of both human and mouse whitened beige adipocytes was induced with 10 *µ*M forskolin for the indicated durations. For all exosome-boosting experiments, Tg-ExoBoost and WT cells were treated with 2 *µ*g/mL doxycycline hyclate (Sigma, D5207) for 72 hours.

### Murine Mouse Models and *In Vivo* Treatment Protocols

All experiments were performed using female mice harvested at 8– 10 weeks of age. Transgenic mice expressing the doxycycline-inducible ExoBooster cassette were generated by the Mouse Genome Modification Core at MaineHealth Institute for Research (MHIR) via pronuclear microinjection of the linearized construct into fertilized C57BL/6J zygotes. The ExoBooster construct includes a Tet-On 3G system and a bidirectional TRE3G promoter, enabling doxycycline-dependent expression of a tricistronic cassette encoding *STEAP3, SDC4*, and *NadB*. Beige adipocytes used for *in vitro* studies were derived from the stromal vascular fraction of inguinal white adipose tissue, as described above. For *in vivo* activation of the Tg-ExoBooster transgene, mice were housed at thermoneutrality (29^*°*^C) for 7 days to normalize adipose tissue whitening between individual mice, then treated with doxycycline (625 mg/kg chow; Inotiv) from days 2–7 and subjected to a 3-day cold challenge (6^*°*^C). For *Rab27a* studies, wild-type, heterozygous, and *Rab27a* null mice were generated as previously described (42). Mice were housed at either room temperature (∼ 22^*°*^C) or cold conditions (6^*°*^C) for 7 days prior to metabolic cage analysis and phenotyping. For CL316,243 studies, mice were housed in metabolic cages and received daily intraperitoneal injections of the *β*_3_-adrenergic receptor agonist CL316,243 (1 mg/kg body weight) for 9 consecutive days to stimulate adipose thermogenesis.

### Metabolic Cage Analysis and Body Composition

Whole-body energy expenditure, respiratory exchange ratio (RER), resting energy expenditure (REE), active energy expenditure (AEE), physical activity (distance walked), food and water intake, and sleep behavior were monitored using the Promethion Core metabolic cage system (Sable Systems International) after a 24-hour acclimation. Body composition was measured using dual-energy X-ray absorptiometry (DXA; Faxitron Bioptics, LLC).

### Exosome Isolation, Quantification and Inhibition

Exosomes were isolated from conditioned medium using the miRCURY Exosome Cell/Urine/CSF Kit (Qiagen) according to the manufacturer’s instructions. Particle size distribution and concentration were measured by nanoparticle tracking analysis (NTA) using the ZetaView instrument (Particle Metrix GmbH). Media was collected from wells seeded with equal numbers of cells and equal volumes, allowing direct comparison of absolute particle counts across conditions. Particle concentrations are reported as particles per mL and shown as absolute values relative to untreated controls. Where indicated, exosome secretion was inhibited by pretreating cells with manumycin A (1 *µ*M, Sigma) or GW4869 (10 *µ*M, Sigma) 2 hours before forskolin treatment, unless otherwise specified.

### qPCR and Western Blotting

Total RNA, including microRNAs, was extracted using the miRNeasy Micro Kit (Qiagen). cDNA was synthesized with qScript cDNA SuperMix (Quanta Biosciences, 95048-100), and quantitative PCR was performed using AzuraView GreenFast qPCR Blue Mix LR (Azura Genomics, AZ-2320) on a Bio-Rad CFX384 real-time PCR system. Gene expression was analyzed using the ΔΔCt method, normalized to *Actb* for mRNA or U6/18S for microRNAs, and expressed relative to vehicle-treated or day 0 controls unless otherwise specified. Melt curve analysis confirmed primer specificity. Protein expression was assessed by Western blot as previously described (46), using *β*-actin or *β*-tubulin as loading controls.

### Heat Production Assay via Microcalorimetry

Heat production from beige adipocytes or freshly isolated adipose tissue was measured using high-resolution isothermal microcalorimetry on the calScreener™ system (Symcel). Equivalent numbers of beige preadipocytes were differentiated in 200 *µ*L of medium (as described above) in Geltrex-coated calWell™ insert plates (Symcel, 1900402). Inserts were transferred to screw-capped titanium vials. The calPlate™ (48-well format) was pre-equilibrated in two thermal zones for 30 minutes, followed by a 15–30 minute stabilization period after final positioning. Continuous heat output was recorded using calView™ software. For *ex vivo* analysis, adipose tissue biopsies were taken from identical locations within the fat pad, placed in calWells containing 200 *µ*L DMEM, and normalized by weight. All measurements were normalized to internal reference wells containing media alone.

### Animal Ethics

All animal studies were conducted in compliance with the National Institutes of Health Guide for the Care and Use of Laboratory Animals and were approved by the Institutional Animal Care and Use Committee (IACUC; protocol #2203) at the MaineHealth Institute for Research. The mice were housed in AAALAC-accredited pathogen-free facilities with *ad libitum* access to food and water. Every effort was made to minimize animal suffering and reduce the number of animals used.

### Data Availability

All data supporting the findings of this study are included in the article and its Supplementary Information. Raw and processed data from RNA and microRNA qPCR screens, microcalorimetry, and metabolic cage studies are available from the corresponding author upon reasonable request.

## Supporting Information Appendix (SI)

### Figures

**Fig. S1:**
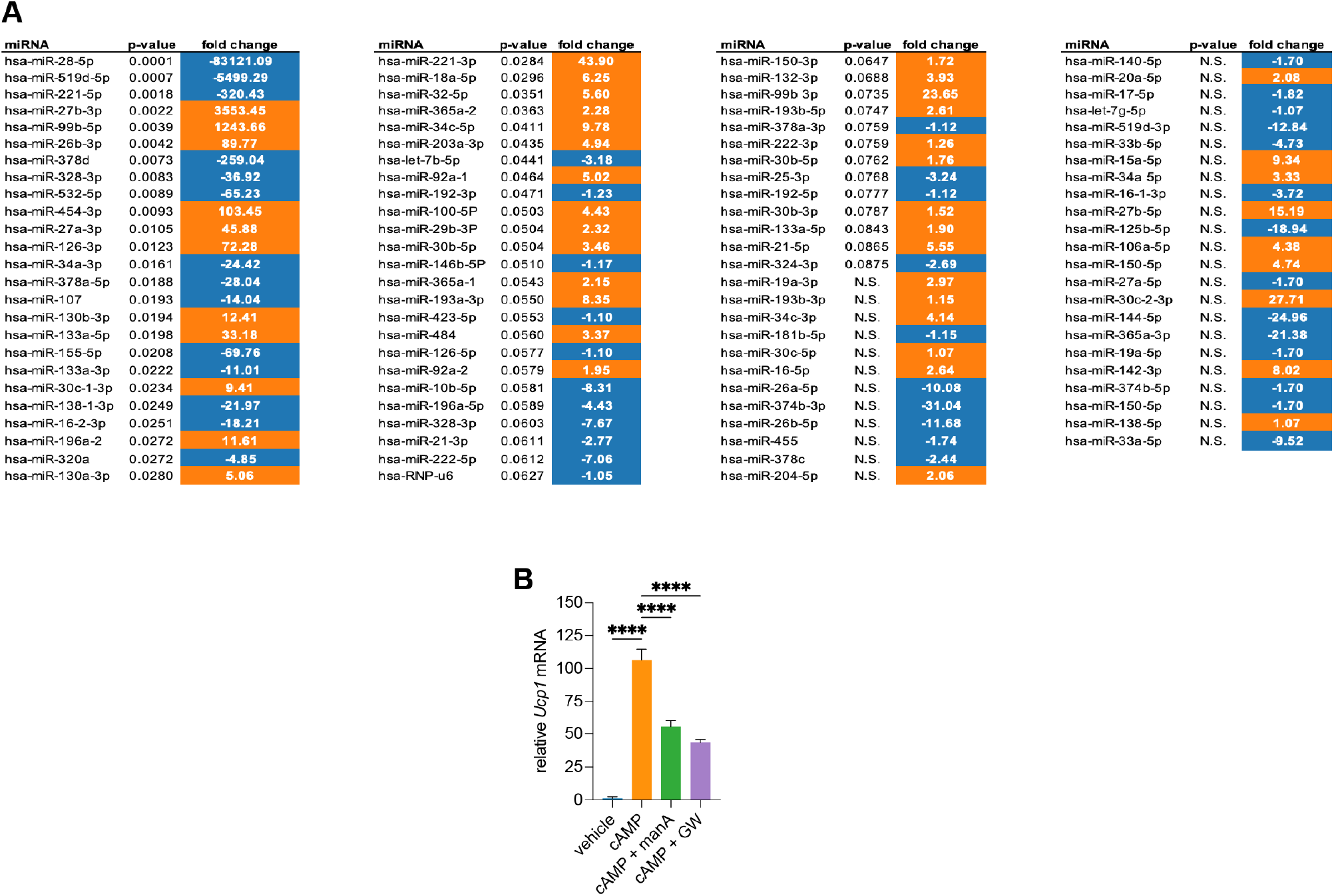
(A) microRNA expression in exosomes from beige adipocytes under whitening versus forskolin-stimulated conditions. Human iPSC-derived beige adipocytes were differentiated for 12 days, then cultured under non-adipogenic (whitening) conditions for 4 days before treatment with forskolin for 6 hours. Exosomes were collected from culture media, and their microRNA content was quantified by targeted qPCR profiling of 98 microRNAs associated with adipogenesis, thermogenesis, and metabolic function. Differential expression analysis was performed using Global Pattern Recognition (GPR) software. This statistical algorithm makes no *a priori* assumptions about normalizers. Inspired by triangulation methods in cartography and astronomy, it iteratively compares each gene’s expression relative to all others in a gene panel to establish a global expression pattern, identifying and ranking significant changes. The plot lists microRNAs with corresponding fold changes sorted by p-values, colored orange for upregulated and blue for downregulated microRNAs following forskolin stimulation compared to whitening conditions. Data represent the mean of *n* = 3 independent experiments. (B) Inhibition of exosome secretion with manumycin A or GW4869 attenuated cAMP-induced upregulation of *Ucp1* mRNA expression in beige adipocytes. Cells were pretreated with manumycin A (1 µM) or GW4869 (10 µM) for 2 hours prior to cAMP stimulation (500 nM) for 6 hours, and *Ucp1* transcript levels were measured by qPCR. These findings indicate that the observed effects are not due to off-target actions of forskolin.

**Fig. S2:**
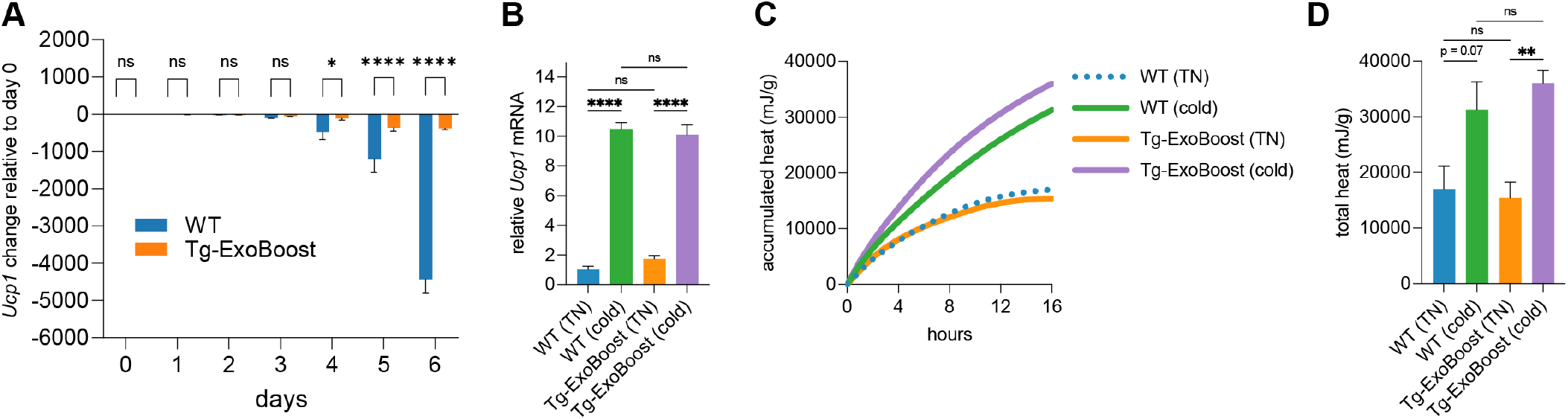
(A) Tg-ExoBooster-positive beige adipocytes maintain *Ucp1* expression during whitening. *Ucp1* mRNA levels in differentiated wild-type (WT) and Tg-ExoBooster beige adipocytes (6 days) cultured in non-adipogenic basal medium for an additional 6 days. Tg-ExoBooster cells maintain higher *Ucp1* expression compared to wild-type. *n* = 4 biological replicates per group; unpaired t-test indicated. (B) Comparable thermogenic responses in brown adipose tissue of WT and Tg-ExoBooster female mice both fed doxycycline diet (3 days) followed by 3 days cold challenge (details in text). RT-qPCR analysis of *Ucp1* expression in brown adipose tissue from wild-type and Tg-ExoBooster mice after doxycycline exposure and cold challenge (*n* = 5). (C, D) Ex vivo microcalorimetry traces showing (C) average heat accumulation over time and (D) total heat output (quantified from C, with error bars) from inguinal fat pads post-cold challenge (*n* = 5). Heat data in C-D normalized to tissue weight in grams (g).

**Fig. S3:**
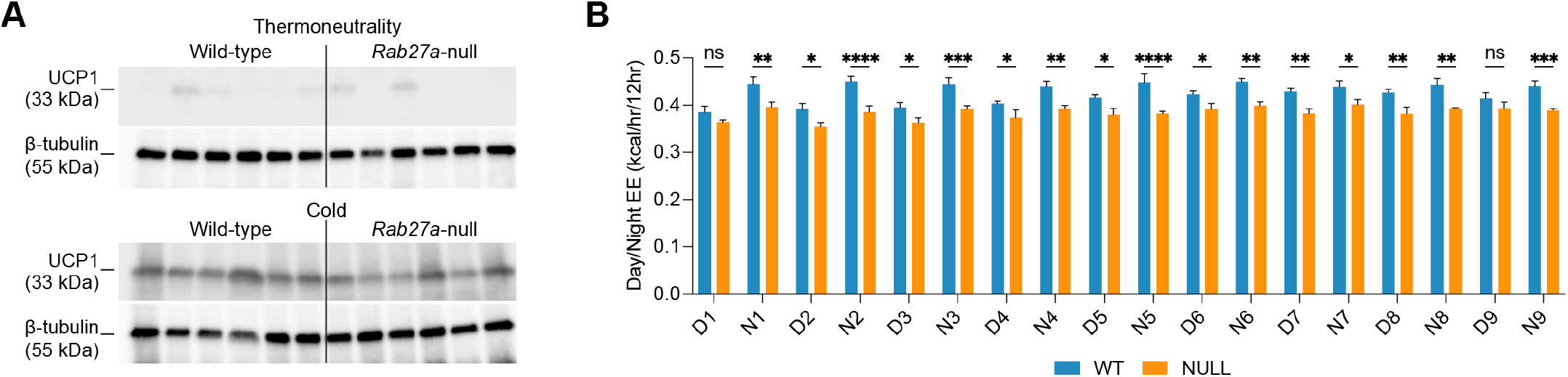
(A) Western blot analysis of UCP1 protein expression in inguinal adipose tissue from wild-type and *Rab27a* null mice following 7 days at thermoneutrality (29 ^*°*^C) and a subsequent 7-day cold exposure. Protein levels were normalized to -tubulin to control for loading. Data correspond to the experiments shown in Fig. 3H. (B) Day-by-day (D) and nighttime energy expenditure profiles (N) in wild-type and *Rab27a* null mice during the 9-day treatment period measured using metabolic cages. Corresponding average energy expenditure across the 9 days is shown in Fig. 4M.

### Tables

**Table S1:** Metabolic and DXA analysis of WT vs *Rab27a* +*/−* mice (RT vs cold).

| Parameter measured | Units | WT | Het | p-value |
| --- | --- | --- | --- | --- |
| Body weight RT (day 7) | g | 20.10 | 21.09 | 0.235 |
| Body weight cold (day 7) | g | 19.45 | 20.27 | 0.3589 |
| Body weight difference (RT vs cold) | % | -3.22 | -3.91 | — |
| Fat mass RT (day 7) | g | 2.53 | 3.20 | **0.0005** |
| Fat mass cold (day 7) | g | 2.35 | 2.48 | 0.4561 |
| Fat mass difference (RT vs cold) | % | -6.93 | -22.40 | — |
| Lean mass RT | g | 13.56 | 13.87 | 0.6407 |
| Lean mass cold | g | 13.12 | 13.65 | 0.4459 |
| Lean mass difference | % | -3.29 | -1.56 | — |
| Daytime energy expenditure RT | kcal/14hr | 0.34 | 0.39 | 0.6782 |
| Daytime energy expenditure cold | kcal/14hr | 0.82 | 0.76 | **0.0037** |
| Nighttime energy expenditure RT | kcal/10hr | 0.47 | 0.46 | 0.8856 |
| Nighttime energy expenditure cold | kcal/10hr | 0.93 | 0.88 | 0.0993 |
| 24h energy expenditure RT | kcal/24hr | 0.43 | 0.44 | 0.5763 |
| 24h energy expenditure cold | kcal/24hr | 0.88 | 0.82 | **0.0200** |
| Daytime RER RT | %/14hr | 0.87 | 0.86 | 0.3416 |
| Daytime RER cold | %/14hr | 0.91 | 0.91 | 0.8286 |
| Nighttime RER RT | %/10hr | 0.92 | 0.93 | 0.4217 |
| Nighttime RER cold | %/10hr | 0.94 | 0.95 | 0.1798 |
| 24h RER RT | %/24hr | 0.90 | 0.89 | 0.9361 |
| 24h RER cold | %/24hr | 0.92 | 0.93 | 0.4739 |
| Daytime REE RT | kcal/30min/14hr | 0.37 | 0.34 | 0.3357 |
| Daytime REE cold | kcal/30min/14hr | 0.73 | 0.68 | **0.0362** |
| Nighttime REE RT | kcal/30min/10hr | 0.37 | 0.34 | 0.3600 |
| Nighttime REE cold | kcal/30min/10hr | 0.79 | 0.73 | **0.0149** |
| 24h REE RT | kcal/30min/24hr | 0.36 | 0.49 | 0.4919 |
| 24h REE cold | kcal/30min/24hr | 0.76 | 0.70 | **0.0114** |
| Daytime AEE RT | kcal/30min/14hr | 0.49 | 0.49 | 0.8010 |
| Nighttime AEE RT | kcal/30min/10hr | 0.52 | 0.54 | 0.5392 |
| 24h AEE RT | kcal/30min/24hr | 0.50 | 0.52 | 0.6286 |
| Daytime AEE cold | kcal/30min/14hr | 0.93 | 0.91 | 0.4956 |
| Nighttime AEE cold | kcal/30min/10hr | 1.02 | 0.95 | 0.1063 |
| 24h AEE cold | kcal/30min/24hr | 0.97 | 0.93 | 0.1952 |
| Food consumed/day RT | g/24hr | 2.01 | 1.44 | 0.1801 |
| Food consumed/day cold | g/24hr | 3.60 | 3.07 | 0.2494 |
| Food consumed/night RT | g/24hr | 2.50 | 2.49 | 0.9746 |
| Food consumed/night cold | g/24hr | 3.32 | 3.36 | 0.9124 |
| 24h food consumed RT | g/24hr | 4.51 | 3.92 | 0.2961 |
| 24h food consumed cold | g/24hr | 6.92 | 6.43 | 0.4146 |
| 24h water consumed/day RT | g/24hr | 3.63 | 3.16 | 0.2452 |
| 24h water consumed/day cold | g/24hr | 3.62 | 3.49 | 0.7817 |
| 24h Y beam breaks (activity) RT | #/24hr | 32314.10 | 31337.20 | 0.8103 |
| 24h Y beam breaks (activity) cold | #/24hr | 33845.00 | 32542.00 | 0.7647 |
| 24h X beam breaks (activity) RT | #/24hr | 22834.10 | 22445.00 | 0.8684 |
| 24h X beam breaks (activity) cold | #/24hr | 23218.20 | 20482.70 | 0.2821 |
| 24h in cage walking meters RT | m/24hr | 316.38 | 229.33 | **0.0462** |
| 24h in cage walking meters cold | m/24hr | 236.00 | 197.67 | 0.3936 |
| 24h in cage walking speed RT | m/s/24hr | 0.03 | 0.03 | 0.9505 |
| 24h in cage walking speed cold | m/s/24hr | 0.03 | 0.03 | 0.3572 |
| 24h Hours of Sleep ($\geq 40$sec) RT | hrs/24hrs | 11.88 | 12.33 | 0.7241 |
| 24h Hours of Sleep ($\geq 40$sec) cold | hrs/24hrs | 11.93 | 11.10 | 0.5494 |

**Table S2:** Metabolic and DXA analysis of WT vs *Rab27a −/−* mice (CL316,243 treated).

| Parameter measured | Units | WT | Null | p-value |
| --- | --- | --- | --- | --- |
| Body weight RT (day 7) | g | 19.05 | 18.78 | 0.6745 |
| Body weight cold (day 7) | g | 19.88 | 19.07 | **0.0323** |
| Body weight difference (RT vs cold) | % | 4.47 | 1.82 | **0.3770** |
| Fat mass RT (day 7) | g | 2.77 | 3.45 | **0.0270** |
| Fat mass cold (day 7) | g | 2.34 | 3.14 | **0.0347** |
| Fat mass difference (RT vs cold) | % | -15.43 | -12.21 | 0.5853 |
| Lean mass pre CL (day 1) | g | 12.56 | 12.16 | 0.4797 |
| Lean mass post CL (day 9) | g | 13.40 | 12.00 | **0.0087** |
| Lean mass change | % | 5.95 | -0.10 | **0.0051** |
| Daytime energy expenditure | kcal/14hr | 0.43 | 0.39 | **0.0062** |
| Nighttime energy expenditure | kcal/10hr | 0.44 | 0.40 | **0.0014** |
| 24h energy expenditure | kcal/24hr | 0.44 | 0.39 | **0.0020** |
| Daytime respiratory exchange ratio | %/14hr | 0.78 | 0.80 | **0.0056** |
| Nighttime respiratory exchange ratio | %/10hr | 0.89 | 0.89 | 0.6171 |
| 24h respiratory exchange ratio | %/24hr | 0.83 | 0.85 | 0.1315 |
| Daytime resting energy expenditure | kcal/30min/14hr | 0.38 | 0.37 | 0.7722 |
| Nighttime resting energy expenditure | kcal/30min/10hr | 0.37 | 0.30 | **0.0326** |
| 24h resting energy expenditure | kcal/30min/24hr | 0.38 | 0.34 | **0.0012** |
| Daytime active energy expenditure | kcal/30min/14hr | 0.49 | 0.48 | 0.6879 |
| Nighttime active energy expenditure | kcal/30min/10hr | 0.50 | 0.44 | **0.0003** |
| 24h active energy expenditure | kcal/30min/24hr | 0.49 | 0.46 | 0.0980 |
| Food consumed/day | g/24hr | 4.63 | 4.01 | 0.2754 |
| Water consumed/day | g/24hr | 3.46 | 2.96 | 0.4984 |
| 24h Y beam breaks (activity) | #/24hr | 23756.17 | 19361.50 | 0.1392 |
| 24h X beam breaks (activity) | #/24hr | 23378.00 | 19429.17 | 0.1742 |
| 24h in cage walking meters | m/24hr | 174.11 | 198.85 | 0.2781 |
| 24h in cage walking speed | m/s/24hr | 0.03 | 0.03 | 0.2015 |
| 24h Hours of Sleep ($\geq 40$sec) | hrs/24hrs | 0.03 | 0.03 | 0.2015 |

